# Aberrant pathogenic GM-CSF^+^ T cells and inflammatory CD14^+^CD16^+^ monocytes in severe pulmonary syndrome patients of a new coronavirus

**DOI:** 10.1101/2020.02.12.945576

**Authors:** Yonggang Zhou, Binqing Fu, Xiaohu Zheng, Dongsheng Wang, Changcheng Zhao, Yingjie qi, Rui Sun, Zhigang Tian, Xiaoling Xu, Haiming Wei

## Abstract

Pathogenic human coronavirus infections, such as severe acute respiratory syndrome CoV (SARS-CoV) and Middle East respiratory syndrome CoV (MERS-CoV), cause high morbidity and mortality ^1,2^. Recently, a severe pneumonia-associated respiratory syndrome caused by a new coronavirus was reported at December 2019 (2019-nCoV) in the city Wuhan, Hubei province, China^3–5^, which was also named as pneumonia-associated respiratory syndrome (PARS)^6^. Up to 9th of February 2020, at least 37, 251 cases have been reported with 812 fatal cases according to the report from China CDC. However, the immune mechanism that potential orchestrated acute mortality from patients of 2019-nCoV is still unknown. Here we show that after the 2019-nCoV infection, CD4^+^T lymphocytes are rapidly activated to become pathogenic T helper (Th) 1 cells and generate GM-CSF etc. The cytokines environment induces inflammatory CD14^+^CD16^+^ monocytes with high expression of IL-6 and accelerates the inflammation. These aberrant and excessive immune cells may enter the pulmonary circulation in huge numbers and play an immune damaging role to causing lung functional disability and quick mortality. Our results demonstrate that excessive non-effective host immune responses by pathogenic T cells and inflammatory monocytes may associate with severe lung pathology. Therefore, we suggest that monoclonal antibody that targets the GM-CSF or interleukin 6 receptor may potentially curb immunopathology caused by 2019-nCoV and consequently win more time for virus clearance.

---

Coronavirus, including SARS and MERS, has caused two large-scale pandemic in the last two decades^1,2^. Although viral evasion of host immune responses and virus-induced cytopathic effects are believed to be critical in disease severity, studies from humans who died of SARS and animal models suggested that an excessive and aberrant host immune response resulting in an exuberant immunopathology and lethal disease^7–9^. Similarly, patients infected with 2019-nCoV, that have been reported recently, have increased plasma concentrations of inflammation related cytokines, including interleukins (IL) 2, 7, and 10, granulocyte-colony stimulating factor (G-CSF), interferon-γ-inducible protein 10 (IP10), monocyte chemoattractant protein 1(MCP1), macrophage inflammatory protein 1 alpha (MIP1A), and tumour necrosis factor α (TNF-α), especially in moribund patients^10^. Importantly, 2019-nCoV infected patients have developed characteristic pulmonary ground glass changes on imaging and lymphocytes decreasing^11,12^. These phenomena suggest severe pulmonary inflammation and cytokine storm also exist in 2019-nCoV infection. At present, symptomatic treatments with organ support to moribund patients are the mainstays of clinical managements. It is urgent to identify the immunopathology mechanism to delay the pulmonary immune injury.

In patients infected with SARS-CoV, it has been reported that the severity of pulmonary immune injury correlated with extensive infiltration of neutrophils and macrophages in the lungs^13,14^, accompanied with increased numbers of neutrophils and monocytes and lower CD8^+^ and CD4^+^ T cell counts in the peripheral blood samples^15–17^. To identify the immune characteristic of patients infected with 2019-nCoV, peripheral blood samples from patients with severe pneumonia were collected for immune analysis. Consistent with previous clinical characteristics reports^18^, these hospitalized patients with confirmed 2019-nCoV infection involved from The First Affiliated Hospital of University of Science and Technology of China commonly have fever symptoms. The patients in intensive care unit (ICU) have significantly decreased concentrations of haemoglobin and albumin, but increased concentrations of C-reactive protein, alanine aminotransferase, aspartate aminotransferase and lactate dehydrogenase (Extended Data Table 1). The number of total leukocytes in peripheral blood had no significant differences between patients of 2019-CoV and healthy controls, whereas the number of lymphocytes decreased significantly in ICU patients. Specifically, monocytes from both ICU and non-ICU patients significantly decreased compared with healthy controls. The number of T cells also significantly decreased from both ICU and non-ICU patients. The CD4^+^ T cells from both patients in ICU and non-ICU decreased remarkably, whereas CD8^+^ T cells decreased more significantly in ICU patients. Other kinds of leukocytes, including granulocyte, B cells and NK cells have no significantly change in numbers between patients of 2019-nCoV and healthy controls (Extended Data Figure. 1).

To demonstrate the status of these aberrant altered T cells, several lymphoid antigens have been analyzed on T cells. These CD4^+^ T cells in patients infected with 2019-nCoV have higher expression of CD69, CD38, and CD44 compared with healthy controls (Fig.1a, b), indicating their activated status. OX40 have been reported to play a major role in promoting clonal expansion and inducing production of several cytokines in T cells^19^. In patients infected with 2019-nCoV, OX40 expression increased remarkably on CD4^+^ T cells, especially in severe ICU patients (Fig.1a, b). CD8^+^T cells in patients infected with 2019-nCoV also showed activated phenotype with higher expression of CD69, CD38 and CD44 (Fig.1c, d). 41BB (CD137; TNFRS9) is an activation-induced co-stimulatory molecule, which is important to priming immune responses of cytotoxic CD8^+^T cells^20^. In ICU patients infected with 2019-nCoV, the expression of 41BB increased significantly compared to healthy controls (Fig.1c, d). It has been reported that co-expression of Tim-3 and PD-1 may represent a subset of T cells with more severe exhaustion in virus infections^21,22^. It is worth noting that much higher percentage of co-expression Tim3^+^PD-1^+^ T subset exist both in CD4^+^ and CD8^+^ T cells from patients of 2019-nCoV (Fig.1e-h), especially in ICU patients, suggesting the exhausted status in T cells in these patients infected 2019-CoV.

**Figure 1.**
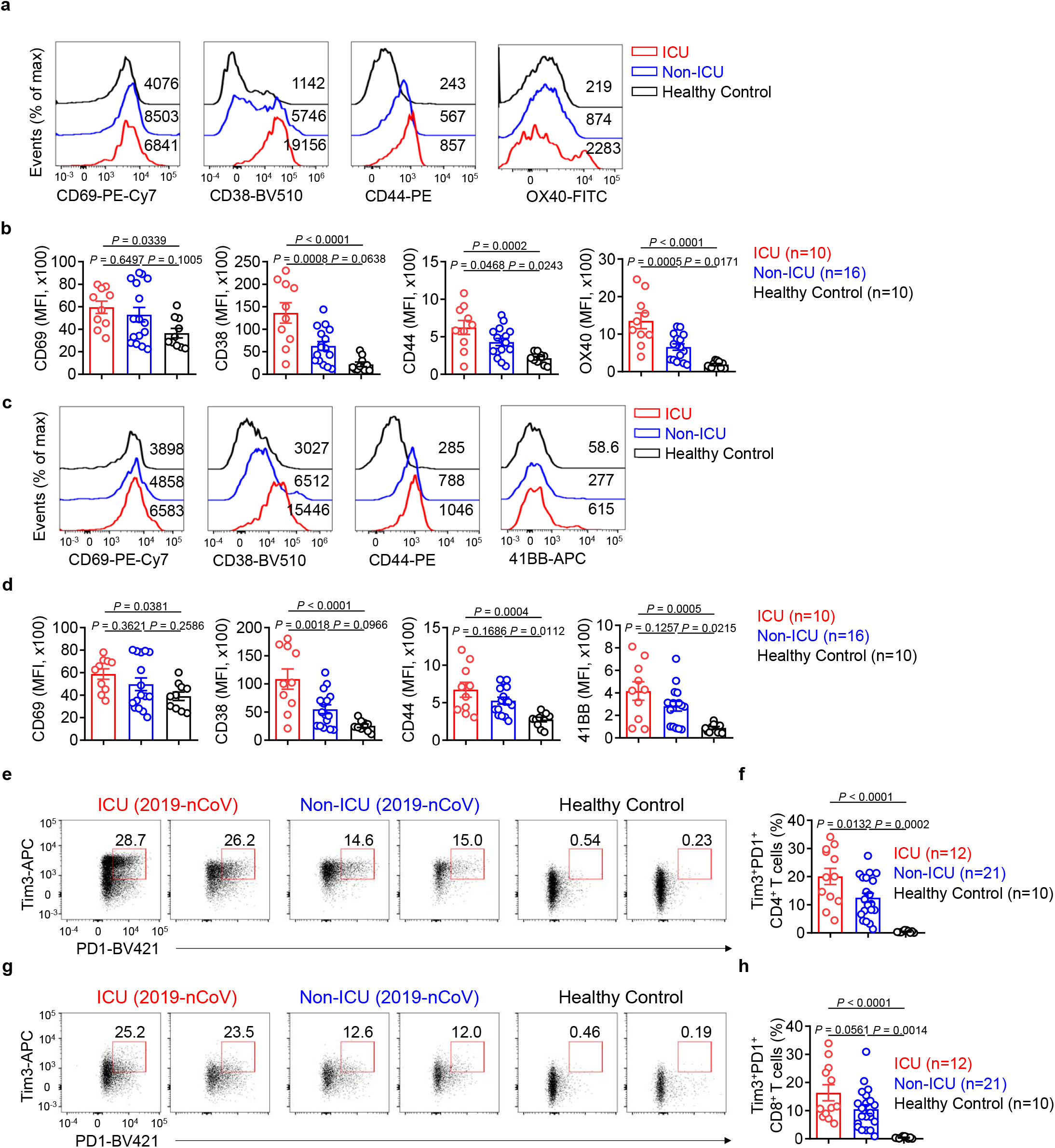
Activated T cells in severe pulmonary syndrome patients of 2019-nCoV. (a, b) Representative density plots and MFI statistics calculated for CD69, CD38, CD44 and OX40 expressions in gated CD3^+^CD4^+^ T cells (Gating strategy showing in Extended Data Figure 2a) isolated from peripheral blood in controls, ICU and non-ICU patients of 2019-nCoV. (c, d) Representative density plots and MFI statistics ed for CD69, CD38, CD44 and 41BB expressions in gated CD45^+^CD3^+^CD8^+^ T cells isolated from peripheral n healthy controls, ICU and non-ICU patients of 2019-nCoV. (e, f) Representative density plots and percentage s calculated for Tim-3 and PD-1 co-expressions in gated CD45^+^CD3^+^CD4^+^ T cells isolated from peripheral blood hy controls, ICU and non-ICU patients of 2019-nCoV. (g, h) Representative density plots and percentage statisticscalculatedforTim-3andPD-1co-expression singated CD45^+^CD3^+^CD8^+^ T cellsisolatedfromperipheralblood in healthycontrols, ICU andnon-ICU patients of 2019-nCoV. Data represent theme an ± SEM.One-wayANOVA. P<0.05 wasconsideredstatisticallysignificant.

To further identify the key pathogenic cytokines and the main source of these cytokines, interferon-γ (IFNγ), TNF-α, granulocyte-macrophage colony-stimulating factor (GM-CSF) and IL-6 have been selected to analyzed through intracellular cytokine staining, for these inflammatory mediators have been proven to be critical as the primary cause of inflammatory cytokine storm in patients infected with SARS-CoV or MERS-CoV^23,24^. Without re-stimulation with PMA or incubation with monensin, high percentage of GM-CSF^+^ and IL-6^+^ expressions could been found in CD4^+^T cells from patients infected with 2019-nCoV in both ICU and non-ICU patients compared to healthy controls (Fig.2a, c). ICU patients with more severe pneumonia showed correlated higher percentage of GM-CSF^+^ and IL-6^+^CD4^+^ T cells (Fig.2a, c). Pathogenic Th1 cells with both IFN-γ and GM-CSF expression have been reported in central nervous system inflammation^25^. Importantly, aberrant pathogenic Th1 cells with co-expressing IFNγ and GM-CSF exist only in ICU patients infected 2019-nCoV, whereas little was found in non-ICU patients and healthy controls, indicating this pathogenic Th1 cells which have correlative evidence from patients with severe disease, play a critical role for hyper-inflammatory responses in 2019-nCoV pathogenesis (Fig.2b, d). Meanwhile, TNF-α were not significant up-regulated in CD4^+^T cells from patients of 2019-nCoV (Extended Data Figure 2a-c). CD8^+^ T cells from ICU patients also showed expression of GM-CSF compared to those from non-ICU patients and healthy controls. IL-6 and TNF-α were not found in CD8^+^ T cells (Extended Data Figure 2d, e). Neither NK cells nor B cells were the secreting source of GM-CSF and IL-6 (Extended Data Figure 2f-i).

**Figure 2.**
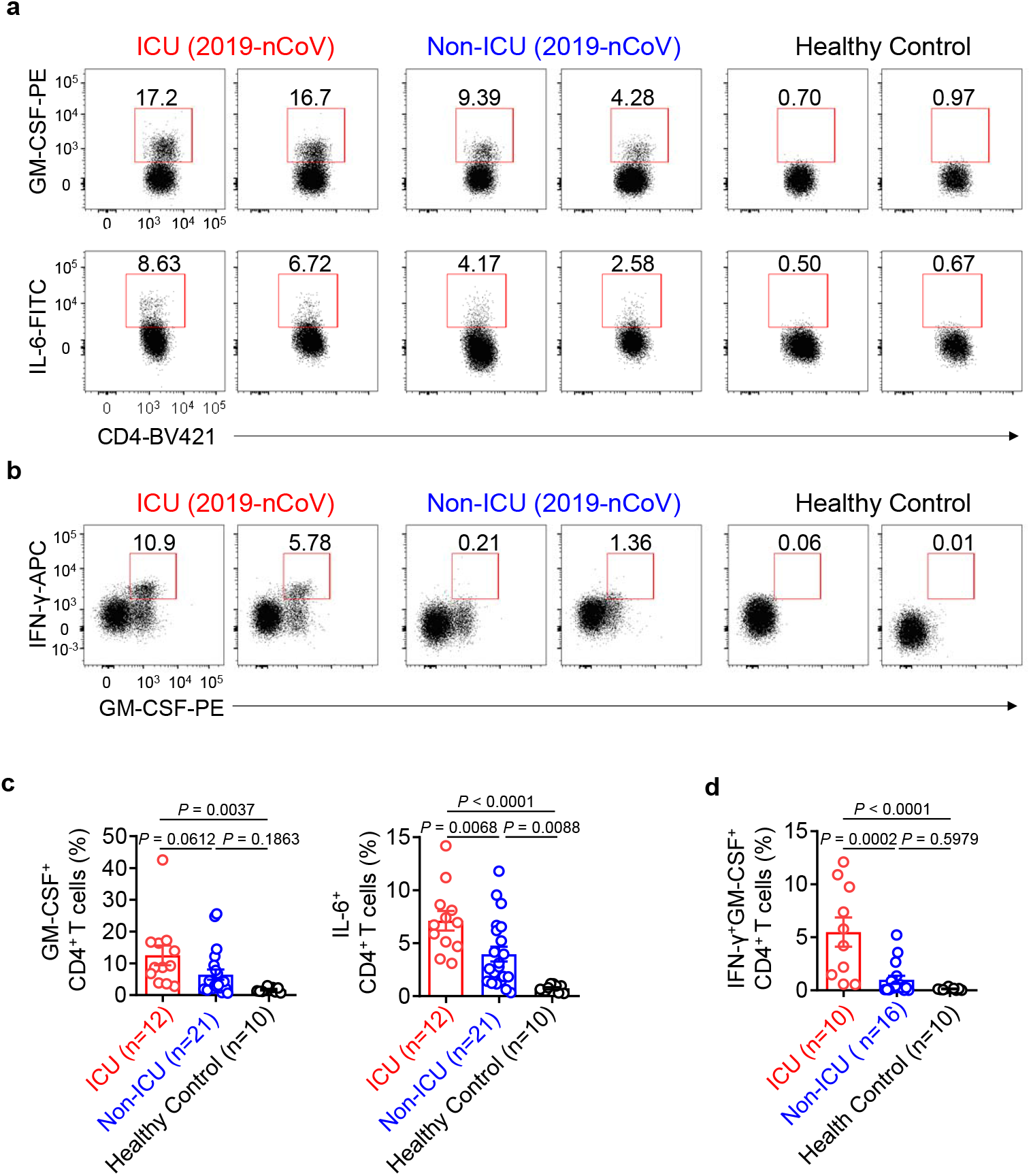
Pathogenic Th1 cells with high expression of GM-CSF in severe pulmonary syndrome patients of 2019-nCoV. Representative density plots showing an analysis of GM-CSF and IL-6 expressions in gated CD45^+^CD3^+^CD4^+^ T cells (Gating strategy showing in Extended Data Figure 1a) isolated from peripheral blood in healthy controls, ICU and non-ICU patients of 2019-nCoV. (b) Representative density plots showing an analysis of co-expression of GM-CSF and IFN-γ in gated CD45^+^CD3^+^CD4^+^ T cells isolated from peripheral blood in healthy controls, ICU and non-ICU patients of 2019-nCoV. (c) Statistics calculated by the percentage of GM-CSF^+^ or IL-6^+^ cells from CD4^+^ T cells. (d) Statistics calculated by the percentage of GM-CSF and IFN-γ co-expressing CD4^+^ T cells. Data represent the mean ± SEM. One-way ANOVA. P<0.05 was considered statistically significant.

GM-CSF has been recently been implicated in the pathogenesis of inflammatory and autoimmune diseases, in a mechanism that controls diverse pathogenic capabilities of inflammatory myeloid cells. Among these myeloid cells, monocyte is the pathogenic GM-CSF responsive cells that require GM-CSF to initiate tissue damage in both mouse and human^26,27^. To identify whether inflammatory monocyte exist in patients infected 2019-nCoV, phenotype and subpopulation of monocytes have been analysis. There was little CD14^+^CD16^+^ inflammatory monocyte subset in healthy controls. By contrast, significant higher percentage of CD14^+^CD16^+^ inflammatory monocyte exist in peripheral blood of patient infected 2019-nCoV. The percentage of CD14^+^CD16^+^ monocyte was much higher in severe pulmonary syndrome patients from ICU (Fig.3a, c). Moreover, these monocyte from patients infected 2019-nCoV also showed capability to secrete GM-CSF. Importantly, significantly higher expression of IL-6 secreted from these inflammatory monocyte especially in ICU patients, which let the cytokine storm even worse (Fig.3b, d). Meanwhile, the number of GM-CSF^+^ monocytes and IL-6^+^ monocytes increased rapidly (Fig.3e), suggesting the potential high risk of inflammatory cytokine storm caused by monocytes that may migrate to the lung and further derive into macrophage or monocyte derived dendritic cells. Thus, in patients infected with 2019-nCoV, GM-CSF potentially links the severe pulmonary syndrome-initiating capacity of pathogenic Th1 cells (GM-CSF^+^IFNγ^+^) with the inflammatory signature of monocytes (CD14^+^CD16^+^ with high expression of IL-6) and their progeny. These activated immune cells may enter the pulmonary circulation in large numbers and played an immune damaging role in severe pulmonary syndrome patients (Fig.4).

**Figure 3.**
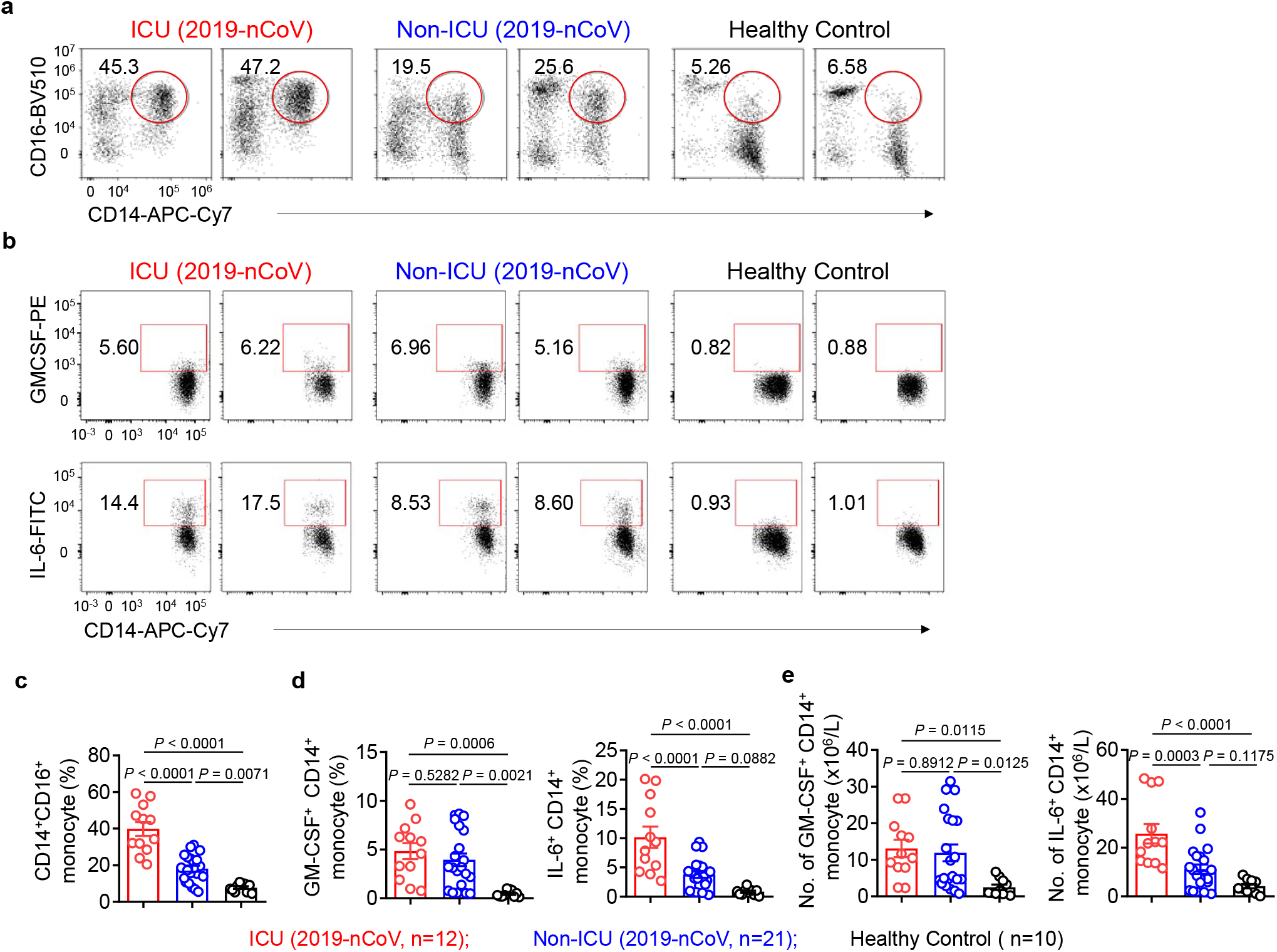
Inflammatory monocytes with high expression of IL-6 in severe pulmonary syndrome patients of 2019-nCoV. (a) Representative density plots showing an analysis of CD14 and CD16 expressions in gated CD45^+^ monocytes strategy showing in Extended Data Figure 1a) isolated from peripheral blood in in healthy controls, ICU and patients of 2019-nCoV. (b) Representative density plots showing an analysis of GM-CSF and IL-6 ons in gated CD45^+^CD14^+^ monocyte cells isolated from peripheral blood in healthy controls, in ICU and patients of 2019-nCoV. (c) Statistics calculated by the percentage of CD14^+^CD16^+^ subsets from monocytes. istics calculated by the percentage of GM-CSF^+^ or IL-6^+^ cells from CD14^+^ monocytes. (e) Statistics d by the cell number of GM-CSF^+^ CD14^+^ or IL-6^+^CD14^+^ monocytes. Data represent the mean ± SEM. ANOVA. P<0.05 was considered statistically significant.

**Figure 4.**
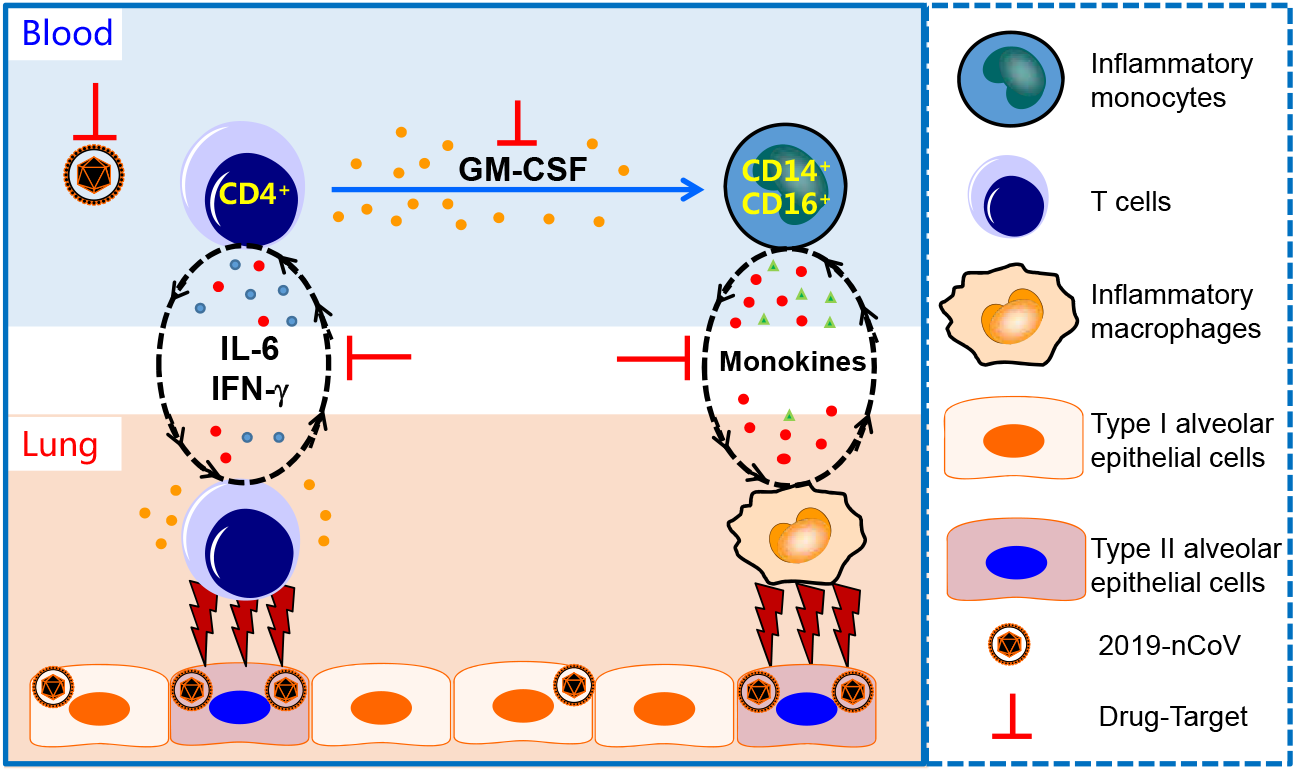
Pathogenic Th1 cells and inflammatory monocytes have positive correlations with severe pulmonary syndrome in patients infected 2019-nCoV. Pathogenic CD4^+^Th1 (GM-CSF^+^IFNγ^+^) cells were rapidly activated to produce GM-CSF and other inflammatory cytokines to form a cascade signature of inflammatory monocytes (CD14^+^CD16^+^ with high expression of IL-6) and their progeny. These activated immune cells may enter the pulmonary circulation in large numbers and played an immune damaging role in severe pulmonary syndrome patients. The monoclonal antibodies that targets the GM-CSF or interleukin 6 receptor may potentially prevent or curb immunopathology caused by 2019-nCoV.

The study provides the detailed immunopathology report on 2019-nCoV, suggesting excessive activated immune response caused by pathogenic GM-CSF^+^ Th1 cells and inflammatory CD14^+^CD16^+^ monocytes may connect pulmonary immunopathology leading to deleterious clinical manifestations and even acute mortality after 2019-nCoV infections. Consistent with the situation with SARS-CoV or MERS-CoV^12,28^, it is remarkable that children always experience mild-moderate clinical illness, elderly individuals exhibit worse outcomes after infection with 2019-nCoV, further indicating that mature excessive immune response towards these pathogenic human coronavirus infections play a key role in inducing severe pulmonary syndrome and even organ failure. However, many urgent questions remain to be answered. Evidence from alveolar washing fluid and lung autopsy from patients infected 2019-nCoV are further needed to verify whether and how these aberrant pathogenic immune cells play a fatal immune damage to cause organ functional disability and mortality. Specific new drugs targeted 2019-nCoV may take long time to evaluate and develop. At this critical moment, several marketed drugs to target cytokine storm and reduce immunopathology could be considered^29^. Blocking inflammatory cytokines may temporarily weaken the anti-infection immunity, yet such strategy is already the lesser of the evils. Other strategies towards blocking the over-activated immune response, such as glucocorticoid treatment showed more side-effect and disappointed outcome towards 2019-CoV^18^. Therefore, we suggest that monoclonal antibody that targets the GM-CSF or interleukin 6 receptor may potentially prevent or curb immunopathology caused by 2019-nCoV and consequently win more time for virus clearance.

## References

1 Drosten, C. et al. Identification of a novel coronavirus in patients with severe acute respiratory syndrome. N Engl J Med 348, 1967–1976, doi:10.1056/NEJMoa030747 (2003).

2 Azhar, E. I., Hui, D. S. C., Memish, Z. A., Drosten, C. & Zumla, A. The Middle East Respiratory Syndrome (MERS). Infect Dis Clin North Am 33, 891–905, doi:10.1016/j.idc.2019.08.001 (2019).

3 Wang, C., Horby, P. W., Hayden, F. G. & Gao, G. F. A novel coronavirus outbreak of global health concern. Lancet, doi:10.1016/S0140-6736(20)30185-9 (2020).

4 Wu, F. et al. A new coronavirus associated with human respiratory disease in China. Nature, doi:10.1038/s41586-020-2008-3 (2020).

5 Zhou, P. et al. A pneumonia outbreak associated with a new coronavirus of probable bat origin. Nature, doi:10.1038/s41586-020-2012-7 (2020).

6 Jiang, S., Xia, S., Ying, T. & Lu, L. A novel coronavirus (2019-nCoV) causing pneumonia-associated respiratory syndrome. Cell Mol Immunol, doi:10.1038/s41423-020-0372-4 (2020).

7 Hui, D. S. C. & Zumla, A. Severe Acute Respiratory Syndrome: Historical, Epidemiologic, and Clinical Features. Infect Dis Clin North Am 33, 869–889, doi:10.1016/j.idc.2019.07.001 (2019).

8 Rockx, B. et al. Early upregulation of acute respiratory distress syndrome-associated cytokines promotes lethal disease in an aged-mouse model of severe acute respiratory syndrome coronavirus infection. J Virol 83, 7062–7074, doi:10.1128/JVI.00127-09 (2009).

9 Smits, S. L. et al. Exacerbated innate host response to SARS-CoV in aged non-human primates. PLoS Pathog 6, e1000756, doi:10.1371/journal.ppat.1000756 (2010).

10 Huang, C. et al. Clinical features of patients infected with 2019 novel coronavirus in Wuhan, China. Lancet, doi:10.1016/S0140-6736(20)30183-5 (2020).

11 Li, G. et al. Coronavirus infections and immune responses. J Med Virol, doi:10.1002/jmv.25685 (2020).

12 Channappanavar, R. & Perlman, S. Pathogenic human coronavirus infections: causes and consequences of cytokine storm and immunopathology. Semin Immunopathol 39, 529–539, doi:10.1007/s00281-017-0629-x (2017).

13 Gu, J. et al. Multiple organ infection and the pathogenesis of SARS. J Exp Med 202, 415–424, doi:10.1084/jem.20050828 (2005).

14 Nicholls, J. M. et al. Lung pathology of fatal severe acute respiratory syndrome. Lancet 361, 1773–1778, doi:10.1016/s0140-6736(03)13413-7 (2003).

15 Cui, W. et al. Expression of lymphocytes and lymphocyte subsets in patients with severe acute respiratory syndrome. Clin Infect Dis 37, 857–859, doi:10.1086/378587 (2003).

16 Li, T. et al. Significant changes of peripheral T lymphocyte subsets in patients with severe acute respiratory syndrome. J Infect Dis 189, 648–651, doi:10.1086/381535 (2004).

17 Wang, Y. H. et al. A cluster of patients with severe acute respiratory syndrome in a chest ward in southern Taiwan. Intensive Care Med 30, 1228–1231, doi:10.1007/s00134-004-2311-8 (2004).

18 Wang, D. et al. Clinical Characteristics of 138 Hospitalized Patients With 2019 Novel Coronavirus-Infected Pneumonia in Wuhan, China. JAMA, doi:10.1001/jama.2020.1585 (2020).

19 Croft, M., So, T., Duan, W. & Soroosh, P. The significance of OX40 and OX40L to T-cell biology and immune disease. Immunol Rev 229, 173–191, doi:10.1111/j.1600-065X.2009.00766.x (2009).

20 Laderach, D., Movassagh, M., Johnson, A., Mittler, R. S. & Galy, A. 4-1BB co-stimulation enhances human CD8(+) T cell priming by augmenting the proliferation and survival of effector CD8(+) T cells. Int Immunol 14, 1155–1167, doi:10.1093/intimm/dxf080 (2002).

21 Khaitan, A. & Unutmaz, D. Revisiting immune exhaustion during HIV infection. Curr HIV/AIDS Rep 8, 4–11, doi:10.1007/s11904-010-0066-0 (2011).

22 Jin, H. T. et al. Cooperation of Tim-3 and PD-1 in CD8 T-cell exhaustion during chronic viral infection. Proc Natl Acad Sci U S A 107, 14733–14738, doi:10.1073/pnas.1009731107 (2010).

23 Drosten, C. et al. Clinical features and virological analysis of a case of Middle East respiratory syndrome coronavirus infection. Lancet Infect Dis 13, 745–751, doi:10.1016/S1473-3099(13)70154-3 (2013).

24 Lew, T. W. et al. Acute respiratory distress syndrome in critically ill patients with severe acute respiratory syndrome. JAMA 290, 374–380, doi:10.1001/jama.290.3.374 (2003).

25 Stienne, C. et al. Foxo3 Transcription Factor Drives Pathogenic T Helper 1 Differentiation by Inducing the Expression of Eomes. Immunity 45, 774–787, doi:10.1016/j.immuni.2016.09.010 (2016).

26 Huang, H. et al. High levels of circulating GM-CSF(+)CD4(+) T cells are predictive of poor outcomes in sepsis patients: a prospective cohort study. Cell Mol Immunol 16, 602–610, doi:10.1038/s41423-018-0164-2 (2019).

27 Croxford, A. L. et al. The Cytokine GM-CSF Drives the Inflammatory Signature of CCR2+ Monocytes and Licenses Autoimmunity. Immunity 43, 502–514, doi:10.1016/j.immuni.2015.08.010 (2015).

28 Assiri, A. et al. Epidemiological, demographic, and clinical characteristics of 47 cases of Middle East respiratory syndrome coronavirus disease from Saudi Arabia: a descriptive study. Lancet Infect Dis 13, 752–761, doi:10.1016/S1473-3099(13)70204-4 (2013).

29 Zumla, A., Hui, D. S., Azhar, E. I., Memish, Z. A. & Maeurer, M. Reducing mortality from 2019-nCoV: host-directed therapies should be an option. The Lancet, doi:10.1016/S0140-6736(20)30305-6.

