## supplemental methods, figures and tables for "Aberrant pathogenic GM-CSF^+^ T cells and inflammatory CD14^+^CD16^+^ monocytes in severe pulmonary syndrome patients of a new coronavirus"

#### Sample collection

Peripheral blood samples of patients infected with 2019-nCoV were collected from The First Affiliated Hospital of University of Science and Technology China (Hefei, Anhui) with the consents from all patients. All of the patients are hospitalized patients with confirmed infection of 2019-nCoV. The diagnosis of pneumonia of unknown cause was first based on clinical characteristics, chest imaging, and ruling out of other pneumonia related bacterial and viral pathogens. Suspected patients were isolated using airborne precautions in the designated hospital. A new coronavirus, which was named 2019-nCoV, was analyzed from lower respiratory tract specimens from these suspected patients. Patients who admitted to the ICU care because they required high-flow nasal cannula or higher-level oxygen support measures to correct hypoxaemia. The detailed patient information has been shown in Extended Data Table 1. Peripheral blood samples from Healthy controls were collected from the First Affiliated Hospital of Anhui Medical University, Hefei, Anhui, China. Ethical approval (No. 2020-XG (H)-005) was obtained from the Ethics Committee from The First Affiliated Hospital of University of Science & Technology of China for emerging infectious diseases.

#### Flow cytometry

Suspensions of lymphocytes were stained for the following human monoclonal antibodies (Extend Data Table 2). Homologous IgGs were used as negative control antibodies. FACS staining was performed according to the manufacturer's instructions. Data from 50,000-80,000 single-cell events were collected using a standard Novocyte3130 flow cytometer (ACEA Biosciences). Intracellular staining of inflammatory cytokines were performed without adding any re-stimulation. The cells were then collected, washed, and blocked according to instructions of eBioscience (Foxp3 / Transcription Factor Staining Buffer Set, eBioscience). The antibodies used for intracellular staining are also shown in Extended Data Table 2.

#### Acknowledgements

This work was supported by the Natural Science Foundation of China (81788101, 81922028), Youth Innovation Promotion Association of Chinese Academy of Sciences (Grant 2019442).

#### Author contributions

Y.Z., B.F. and X.Z performed the experiments, analyzed and interpreted the data. D.W., C. Z., and Y.Q. helped to collect samples and information from patients. R.S. established techniques of FACS and interpreted the data. Z.T. provided strategic planning and interpreted some data. X.X supervised the clinical treatment of patients of 2019-CoV and helped with data interpretation. H.W. supervised the project, provided crucial ideas, and assisted with data interpretation. B.F. wrote the manuscript with H.W.

#### Competing interests

The authors declare no competing interests.

#### Additional information

**Extended Data Figure1.**

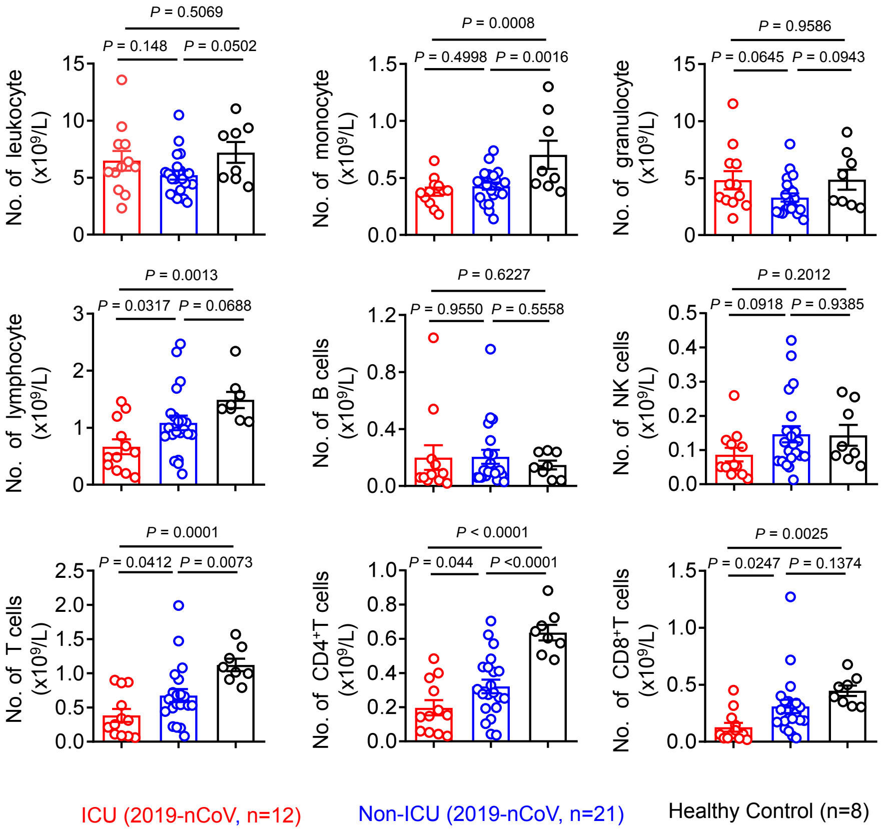


**Extended Data Figure 1. The altered composition of white blood cells in 2019-nCoV patients.** Quantification of leukocyte, monocyte, granulocyte, lymphocyte, NK cells, B cells, CD4+ and CD8+T cells from flow cytometry analysis in 2019-nCoV. Each dot represents one samples. Data represent the mean ± SEM. Data were analyzed by one-way ANOVA. $P<0.05$ was considered statistically significant.

**Extended Data Figure 2.**

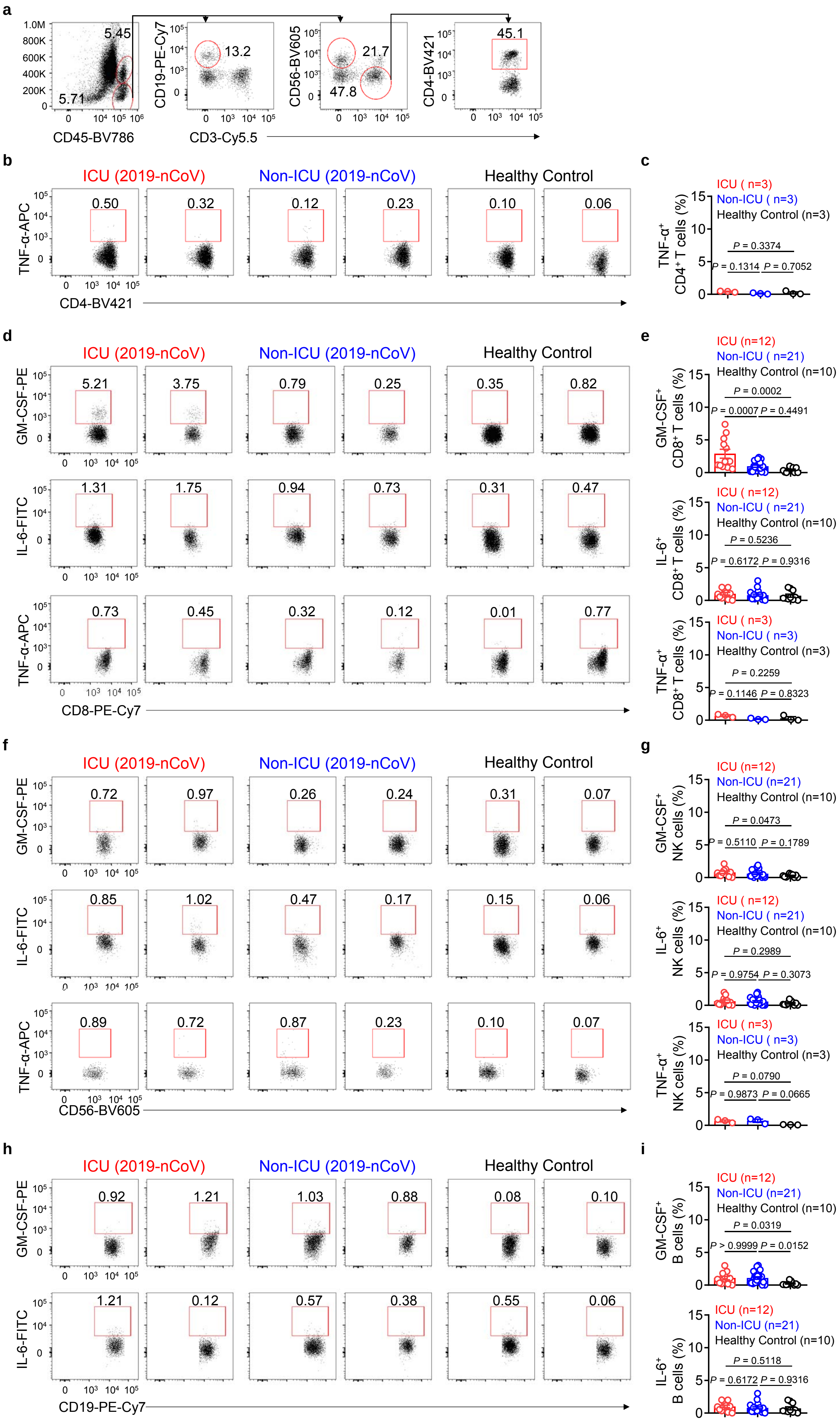

**Extended Data Figure 2. The inflammatory cytokines expression characteristic in T cells, NK cells and B cells from 2019-nCoV patients.**

(a)Gating strategy for monocyte, lymphocytes, $CD3^{+}$T cells, $CD19^{+}$B cells, $CD56^{+}$NK cells, $CD4^{+}$T cells and $CD8^{+}$T cells. (b, c) Representative density plots and percentage statistics calculated for TNF-$\alpha$ expressions in gated $CD45^{+}CD3^{+}CD4^{+}$ T cells isolated from peripheral blood in healthy controls, ICU and non-ICU patients of 2019-nCoV. (d, e) Representative density plots and percentage statistics calculated for GM-CSF, IL-6 and TNF-$\alpha$ expressions in gated $CD45^{+}CD3^{+}CD8^{+}$ T cells isolated from peripheral blood in healthy controls, ICU and non-ICU patients of 2019-nCoV. (f, g) Representative density plots and percentage statistics calculated for GM-CSF, IL-6 and TNF-$\alpha$ expressions in gated $CD45^{+}CD3^{-}CD56^{+}$ NK cells isolated from peripheral blood in healthy controls, ICU and non-ICU patients of 2019-nCoV. (h, i) Representative density plots and percentage statistics calculated for GM-CSF and IL-6 expressions in gated $CD45^{+}CD3^{+}CD19^{+}$ B cells isolated from peripheral blood in healthy controls, ICU and non-ICU patients of 2019-nCoV. Each dot represents one samples. Data represent the mean $\pm$ SEM. Data were analyzed by one-way ANOVA. $P<0.05$ was considered statistically significant.

**Extended Data Table 1. Baseline characteristics and laboratory findings of patients infected with 2019-nCoV in this study**

| | ICU care (n=12) | No ICU care (n=21) | P value |
|---|---|---|---|
| Age, median(IQR), years | 50(29-68) | 41.8 (22-76) | 0.12 |
| Gender(M/F), No.(%) | 9(75%)/3(25%) | 13(61.9%)/8(38.1%) | 0.46 |
| Symptomes, fever, No.(%) | 12(100%) | 21 (100%) | |
| White blood cell count, $X10^9/L$, Median (IQR) | 6.29 (2.32-13.58) | 5.17 (2.82-10.49) | 0.23 |
| Neutrophil count, $X10^9/L$, Median (IQR) | 4.98 (1.79-12.24) | 3.36 (1.37-10.03) | 0.10 |
| Monocyte count, $X10^9/L$, Median (IQR) | 0.36 (0.11-0.52) | 0.33 (0.17-0.65) | 0.63 |
| Lymphocyte count, $X10^9/L$, Median (IQR) | 0.86 (0.32-1.69) | 1.33 (0.28-2.86) | 0.04 |
| Haemoglobin (g/l), Median (IQR) | 125.42 (97-144) | 145.24 111-162) | 0.002 |
| Platelet count, $X10^9/L$, Median (IQR) | 194.58 (72-307) | 184.88 (131-292) | 0.64 |
| C-reactive protein (mg/L), Median (IQR) | 75.13 (8.7-175.2) | 13.15 (0.5-91.1) | 0.0002 |
| Activated partial thromboplastin time, s, Median (IQR) | 36.32 (27.9-42.9) | 41.67 (24.4-67.8) | 0.09 |
| Prothrombin time, s, Median (IQR) | 14.23 (12.7-15.3) | 21.08 (12.1-142) | 0.46 |
| D-dimer, mg/L, Median (IQR) | 0.61 (0.08-3.93) | 0.17 (0.02-0.52) | 0.09 |
| Albumin, g/L, Median (IQR) | 37.08 (29.9-47.6) | 44.09 (37.3-53.6) | 0.0015 |
| Alanine aminotransferase, U/L, Median (IQR) | 43 (16-95) | 25 (12-15) | 0.0021 |
| Aspartate aminotransferase, U/L, Median (IQR) | 40.25 (19-70) | 24.71 (13-44) | 0.0014 |
| Total bilirubin, mmol/L, Median (IQR) | 21.27 (7.9-53.8) | 18.05 (8.8-37.5) | 0.45 |
| Potassium, mmol/L, Median (IQR) | 3.85 (3.03-4.71) | 4.07 (3.43-4.81) | 0.22 |
| Sodium, mmol/L, Median (IQR) | 136.75 (133-142) | 138.88 (131-146) | 0.12 |
| Creatinine, μmol/L, Median (IQR) | 72.75 (54-96) | 74.47 (54-101) | 0.76 |
| Creatine kinase, IU/L, Median (IQR) | 220.46 (49.6-1011.4) | 108.39 (31.3-273.9) | 0.19 |
| Lactate dehydrogenase, U/L, Median (IQR) | 325.82 (189-513) | 219.63 (117-338) | 0.0021 |
| Hypersensitive troponin I, pg/mL, Median (IQR) | 0.08 (0.03-0.12) | 0.07 (0.04-0.1) | 0.11 |
| Procalcitonin, ng/mL, Median (IQR) | 0.19 (0.1-0.37) | 0.19 (0.1-0.22) | 0.47 |

Abbreviations: ICU, intensive care unit; IQR, interquartile range; P values indicate differences between ICU and non-ICU patients.

P<0.05 was considered statistically significant. P values comparing ICU care and no ICU care are from Fisher's exact test or unpaired T test.

### Extended Data Table 2 Antibody list

| REAGENT | SOURCE | IDENTIFIER |
|---|---|---|
| Anti-human CD3 PerCP-CY5.5 | Biolegend | Cat#300328 |
| Anti-human CD14 APC-CY7 | BD Pharmingen | Cat#557831 |
| Anti-human CD16 BV510 | Biolegend | Cat#563830 |
| Anti-human CD56 BV605 | Biolegend | Cat#362538 |
| Anti-human CD45 BV786 | Biolegend | Cat#563716 |
| Anti-human CD4 BV421 | Biolegend | Cat#562424 |
| Anti-human CD8 PE-CY7 | BD Pharmingen | Cat#557746 |
| Anti-human CD19 PE-CY7 | BD Pharmingen | Cat#557835 |
| Anti-human IL6 FITC | BD Pharmingen | Cat#340526 |
| Anti-human GM-CSF PE | BD Pharmingen | Cat#554507 |
| Anti-human IFN-γ APC | BD Pharmingen | Cat#551385 |
| Anti-human TNFα APC | BD Pharmingen | Cat#551384 |
| Anti-human CD69 PE-CY7 | BD Pharmingen | Cat#557745 |
| Anti-human OX40 FITC | BD Pharmingen | Cat#555837 |
| Anti-human 41BB APC | BD Pharmingen | Cat#550890 |
| Anti-human CD44 PE | BD Pharmingen | Cat#555479 |
| Anti-human CD38 BV510 | BD Pharmingen | Cat#563251 |
| Anti-human PD1 BV421 | BD Pharmingen | Cat#562516 |
| Anti-human Tim3 APC | BD Pharmingen | Cat#565558 |
| Mouse IgG2a, κ FITC | BD Pharmingen | Cat#555573 |
| Mouse IgG2b, κ FITC | BD Pharmingen | Cat#555742 |
| Mouse IgG1, κ PE | BD Pharmingen | Cat#555749 |
| Mouse IgG2a, κ PE | BD Pharmingen | Cat#555574 |
| Mouse IgG2a APC | BD Pharmingen | Cat#555576 |
